# Electrostatic modulation of signaling at cell membrane: Waveform- and time-dependent electric control of ERK dynamics

**DOI:** 10.1101/2023.08.31.555453

**Authors:** Minxi Hu, Houpu Li, Kan Zhu, Liang Guo, Min Zhao, Huiwang Zhan, Peter N. Devreotes, Quan Qing

## Abstract

Different exogenous electric fields (EF) can guide cell migration, disrupt proliferation, and program cell development. Studies have shown that many of these processes were initiated at the cell membrane, but the mechanism has been unclear, especially for conventionally non-excitable cells. In this study, we focus on the electrostatic aspects of EF coupling with the cell membrane by eliminating Faradaic processes with dielectric-coated microelectrodes, and show that the ERK signaling pathway of epithelial cells (MCF10A) can be both inhibited and activated by AC EF with different amplitude thresholds, peaking times and refractory periods. Interestingly, the ERK responses were sensitive to the waveform and timing of EF stimulation pulses, depicting the characteristics of electrostatic and dissipative interactions. Blocker tests and correlated changes of active Ras on the cell membrane with ERK signals indicated that both EGFR and Ras were involved in the rich ERK dynamics induced by EF. We propose that the frequency-dependent dielectric relaxation process could be an important mechanism to couple EF energy to the cell membrane region and modulate membrane protein-initiated signaling pathways, which can be further explored to precisely control cell behavior and fate with high temporal and spatial resolution.

## INTRODUCTION

Electric field (EF) has long been exploited to facilitate cell functional development, proliferation, and migration ^1, 2, 3, 4^. There are many biomedical and clinical applications of EF, such as tissue engineering and regeneration ^5, 6^, wound healing^7, 8, 9, 10^, drug delivery ^11, 12, 13^, and cancer treatment and stem cell therapy ^14, 15, 16^. However the quantitative understanding of how EF couples with cells has been unsatisfactory ^2, 17, 18^. EF could bring many biophysical effects on cells, including disruption of structural water, electroosmotic flow, mechanosensation, asymmetric ion flow/opening of voltage-gated channels, and redistribution of membrane components and lipid rafts ^2^. In addition, electrochemical processes associated with applying EF can bring changes in pH and reactive oxidative species (ROS), which are also shown to promote cell responses ^19^. It has been widely recognized that a variety of EFs, including direct-current (DC) ^3, 8^, low-frequency AC EF (less than several hundred Hz) ^9, 20, 21^, fast nanosecond pulses ^22, 23^, and high-frequency radiations (several GHz) ^9^, can cause changes in cytosolic Ca^2+^ levels, and activate p38, c-Jun N-terminal kinase (JNK) and extracellular-signal-regulated kinase (ERK) signaling pathways ^9, 19, 22, 23, 24^, which regulate a lot of stimulated cellular processes and play important roles in cell survival ^25, 26, 27^, motility ^28, 29^, differentiation and proliferation ^30, 31, 32^. However, since many physical or secondary chemical factors may contribute to the final cell response, and during these studies the EF was typically applied to cells for long hours before quantification by population average, the nature of the coupling mechanism between EF and intracellular signaling pathways, particularly in conventionally non-excitable/non-electrogenic cells, remains obscure or controversial at the molecular level.

Early in the 1990s, it was proposed that the constituents of the cell membrane would be much better detectors of weak EF than isolated molecules in solution ^33, 34^. If a signal averaging occurs through field-induced variation in the catalytic activity of a membrane protein, such as a simple Michaelis-Menten-type enzyme embedded in a membrane, cell response could be frequency-dependent ^33, 35^. To date, only the Na-K pump system, which is known to be sensitive to the potential and chemical gradient across the membrane, has demonstrated this behavior ^36, 37, 38^. In these cases, EF can thermodynamically offset the energy states of the ion pumps, causing the pumping rate to eventually synchronize with the period of the external driving field. In addition, low-intensity EF at an intermediate frequency, named tumor treating fields (TTFields), have been shown to block cell division and interfere with organelle assembly, which is attributed to ionic condensation waves around microtubules and dielectrophoretic effects on the dipole moments of microtubules ^39, 40, 41^. Theoretical models have suggested that the energy of EF below 10MHz should be mainly focused on the membrane region due to its high impedance in this frequency range ^42^. However, it remains unclear if it is possible to electrostatically modulate the membrane enzymatic activities that are generally considered *not* sensitive to the electric field, in other words, in conventional non-excitable cells.

Interestingly, some of these membrane receptors, such as epidermal growth factor receptor (EGFR), have been shown to play an important role in EF-induced activation of ERK signaling pathways ^9, 19, 22, 23, 43^. For example, Wolf-Goldberg et al. showed that low-frequency unipolar EF pulses can cause ligand-independent activation of EGFR-Ras-ERK signaling pathway ^19^, which was attributed to the electrochemical elevation of H^+^ and ROS concentrations. On the other hand, in the recent studies from Guo et al., where net ionic current and Faradaic processes have been deliberately avoided by using high-k dielectric passivated microelectrodes and low amplitude symmetrical bipolar EF pulses, it is observed that EF could still locally induce ligand-independent EGFR phosphorylation and ERK activation ^43^. In addition, it has been shown that EF can be used to precisely control synchronized ERK activation patterns to regulate cell fate ^44^. These studies showed the great potential of using EF to modulate the ERK signaling pathway, whereas it is also clear that the coupling mechanisms between exogenous EF and membrane protein-initiated signaling pathways need further investigation.

Here we present a systematic evaluation of the electrostatic aspect of the coupling between EF and the EGFR-Ras-ERK signaling pathway and focus on the immediate response from cells within 0-60 minutes. We observed bipolar ERK response to AC EF stimulation and investigated how different waveforms and timing changed the ERK dynamics. With additional inhibitor tests and quantification of Ras protein in response to EF stimulation, the possible mechanism of how EF energy impacts the enzymatic activities of membrane proteins is discussed.

## METHODS

### Overall experiment design

The EGFR-Ras-ERK signaling pathway starts with two important steps at the membrane, where many moving charges and dipoles of the proteins can interact (Figure 1A). First, after binding with EGF at its extracellular domain, the intracellular C-terminal of EGFR gets phosphorylated, where the adaptor protein (Grb2) and Ras GTP exchange factor (Sos) will dock to facilitate the activation of Ras-GTP ^45^. Second, the Ras-GTP, which is anchored to the membrane through its hydrophobic lipidated C-terminal, forms a complex that binds and activates Raf kinase, which then passes downstream signaling to MEK and ERK in the cytosol.

**Figure 1.**
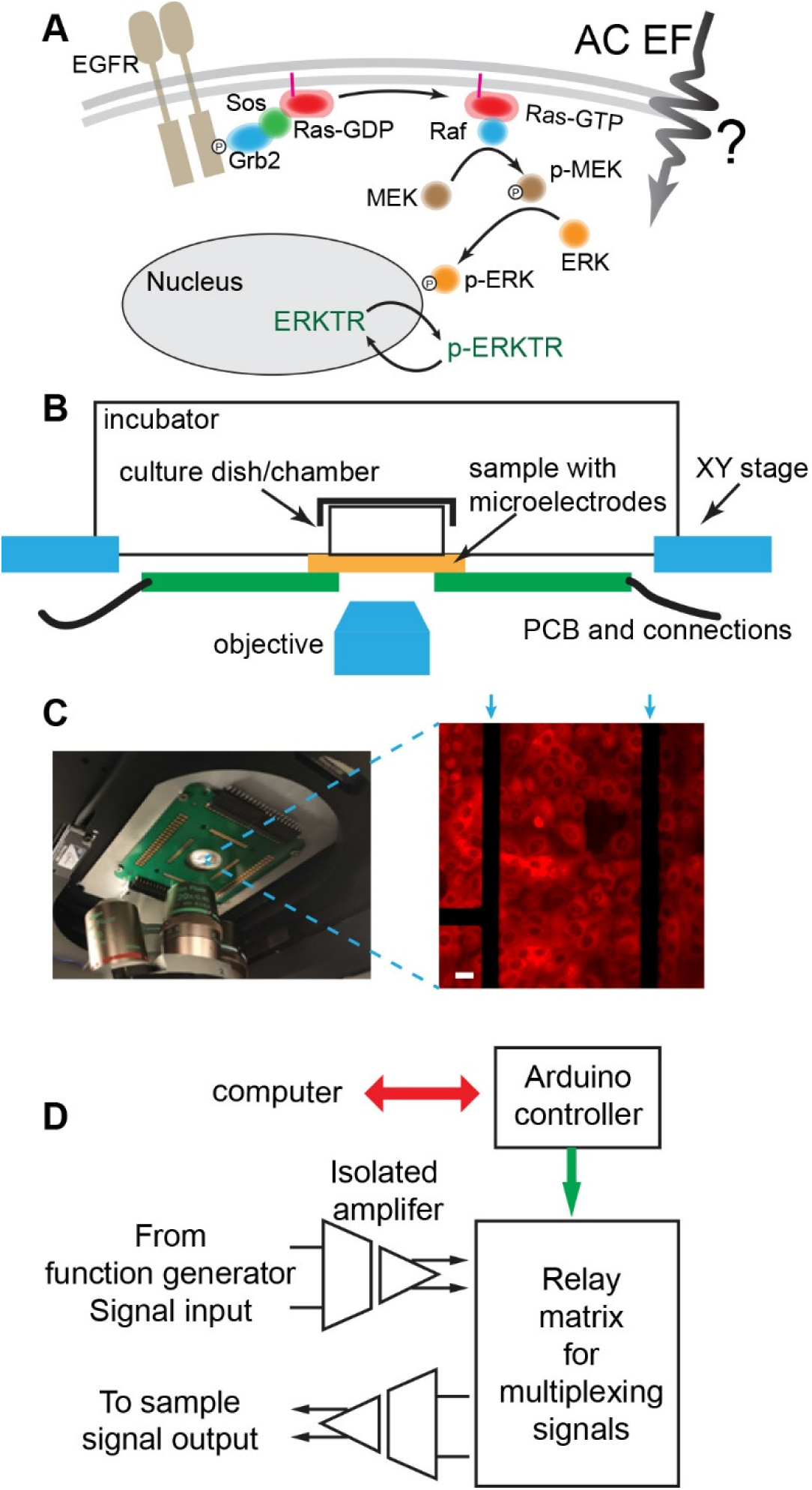
Schematics and experiment setup of electrostatic modulation of EGFR-Ras-ERK signaling pathway. **(A)** The signaling cascade of the EGFR-Ras-ERK pathway. **(B)** Schematics of the on-stage incubation chamber with microelectrode chip integrated for the EF perturbation. **(C)** Left: Picture of the PCB board mounted on the inverted microscope stage with microelectrode chip in the center. Right: Typical fluorescence image of MCF10A cells cultured on top of the electrodes. The dark nucleic area shows that ERKTR is phosphorylated and transported into cytosol due to the high ERK level. The arrows mark the position of the microelectrodes. Cells within 100 µm distance from the microelectrodes are used to calculate the ERKTR ratio. Scale bar: 10 µm. **(D)** Diagram of the customized electronic controls to deliver electric pulses to the sample. An Arduino controller is programmed to switch the relay matrix and direct signals from a function generator to selected microelectrodes. The signals are transmitted through isolated amplifiers so that the microelectrodes do not share ground with any other systems.

We will send bipolar electrical pulses in the frequency range of tens of kHz (with zero DC component) of the same amplitude but different waveforms and timing to live cells through microelectrodes coated with an ultra-thin high-k dielectric layer, where only capacitively coupled AC signals are allowed to pass such that there are no electrochemical complications to the cell environment. Due to the heterogeneous impedance distribution across the cell, EF is mainly induced in the cell membrane and excluded from the highly conductive cytoplasm and culture medium ^42^. We monitor the ERK dynamics of individual cells as a response to the EF stimulation with the ERK transportation reporter (ERKTR) ^43, 46^, which resides in the nucleic region when the ERK level is low, and will be phosphorylated and transported out to the cytosol when the ERK level becomes high. The ratio of the fluorescence intensity of ERKTR in the cytosol and the nucleic (Fc/Fn) can be used to quantify the ERK level with a better than 1-minute time resolution.

### Microscope and EF stimulation setup

A customized on-stage incubator system has been built for the stimulation and imaging experiments (Figure 1B). Specifically, the transparent microelectrode chip is bonded on a printed circuit board (PCB) and attached to the bottom side of an on-stage incubator. A 35-mm diameter petri dish with a 13-mm hole is sealed on top of the chip, serving as the culture chamber. The temperature, humidity, and CO_2_ concentration of the incubator are regulated to support cells cultured on top of the chip. A picture of the mounted chamber with the chip and PCB and the typical fluorescence image of ERKTR being transported out of the nucleic region at a high ERK level are shown in Figure 1C. In addition, we have developed a multiplexing system so that the electrical pulses can be sent to selected electrodes through a relay matrix programmed by an Arduino controller (Figure 1D). Isolation amplifiers (Analog Devices, AD215) are used to transmit the signals, which physically isolates the microelectrodes from all other instruments to cut off any current leakage path or ground loops. In addition, the microelectrodes are passivated by a thin insulation layer of HfO_2_ deposited by atomic layer deposition (ALD) process. These designs will eliminate electrochemical reactions at the interface of the microelectrode and the medium such that we can focus on the pure effect of the locally induced electric field. Details of the experiment procedures are described below.

### Cell culture

MCF10A in co-expression of EKAR3 and ERKTR (MCF10A-EKAR3-ERKTR) was kindly provided by Dr. John Albeck (see Acknowledgements). The cell transfection method is described in reference ^47^. Briefly, co-transfection of pPBJ-EKAR3-nes and the pCMV-hyPBase transposase vector was used to make cells stably expressing EKAR3, then MCF10A-EKAR3 cells were infected by retroviral particles carrying ERKTR-mCherry.

MCF10A in co-expression of Raf-RBD-GFP and Lyn11-FRB-CFP was kindly provided by Dr. Devreotes’ Lab. The cell transfection method is described in reference ^48^. Briefly, transient transfection was conducted using Lipofectamine 3000 reagent (ThermoFisher Scientific) according to the manufacturer’s protocol. Stable transfected MCF10A cells were selected and/or maintained in the culture medium with 2 µg/ml Puromycin (ThermoFisher Scientific) and 2 mg/ml Zeocin (ThermoFisher Scientific).

Both MCF10A cell lines were cultured in Dulbecco’s modified Eagle’s medium (DMEM)/F-12 (Gibco) supplemented with 5% horse serum (Gibco), 0.5 mg/ml hydrocortisone (Sigma-Aldrich), 10 mg/ml human insulin (Gibco), 20 ng/ml EGF (Life Technologies), 100 ng/ml cholera toxin (Sigma-Aldrich), 50 U/ml penicillin and 50 U/ml streptomycin (Gibco). Cells were incubated in a humidified atmosphere containing 5% CO_2_ at 37 °C. After plating on the microelectrode chip, cells were incubated for 12 hours before the container is mounted on the microscope for the EF stimulation experiment.

### Fabrication and assembly of microelectrode chip

The fabrication of microelectrodes was performed on #1 (170 µm thick), 24 × 40 mm microscope cover glass (Thermo Scientific). First, the cover glass was treated with piranha solution, a mixture of sulfuric acid (Sigma-Aldrich) and hydrogen peroxide (GFS Chemicals), to clean organic residues and hydroxylate the surface. Then a layer of 740 nm silicon oxide was deposited on the glass surface using plasma-enhanced chemical vapor deposition (PECVD) (Oxford PLASMALAB 100 PECVD). Standard photolithography and thermal evaporation were then used to pattern the Cr/Au (1/25 nm) microelectrodes on the glass surface. The microelectrodes were shaped as pairs of parallel bars (10 µm wide and 350 µm long) with a gap distance of 80-110 µm, which were connected to 200 µm × 200 µm bonding pads placed at the edges of the chip by thin metal lines. Next, the surface was sputtered with 1 nm Cr (Lesker PVD75) to promote better adhesion before a uniform 10-100 nm HfO_2_ layer was deposited with atomic layer deposition (ALD) (Cambridge Savannah ALD system). Last, an insulating layer of 500 nm SU-8 (MicroChem Inc., SU-8 2000.5) was patterned by standard photolithography such that only the parallel bars of the microelectrodes were exposed for making contact with the solution. The microelectrode chip was then glued on a customized printed circuit board (PCB) with a center hole for optical access, and wire bonded (West-Bond 7KE). Afterward, a 35 × 10 mm petri dish with a 13 mm diameter hole on the bottom was sealed on the chip using Kwik-Sil silicone elastomer (World Precision Instruments) to serve as a cell chamber and medium reservoir.

### Live cell imaging

A customized on-stage incubator with regulated temperature (37 °C), humidity (95%), and CO_2_ concentration (5%), was used to maintain cell culture conditions on the microscope during the imaging process. Time-lapse images were acquired from a Nikon Eclipse Ti-U microscope equipped with a PCO.Edge 4.2 sCMOS camera and a SOLA SE II 365 light engine. Cell images were taken using a 20X CFI S Plan Fluor Ph1 objective with 0.45 numerical aperture and recorded via NIS-Elements software. ET-mCherry and ET-EYFP (49008 and 49003, Chroma Technology Corp) filter sets were used in mCherry and YFP channels, respectively. The images were taken every 30 seconds.

Before cell plating, the assembled microelectrode chip with PCB was attached to the open bottom of the incubator. The chip surface and the cell chamber were sterilized with 75% ethanol and then treated with FNC coating mix (AthenaES) to increase cell attachment. MCF10A-EKAR3-ERKTR cells maintained in culture flasks were dissociated into single cells with 0.25% trypsin-EDTA (Gibco) and seeded in the cell chamber with a density of ∼0.15 million cells/cm^2^. DMEM/F12 without phenol red (Gibco) was used in all cell imaging experiments. In experiments where the starting ERK is maintained at a normal level, the EGF-containing recording medium was supplemented with 5% horse serum, 0.5 mg/ml hydrocortisone, 10 mg/ml human insulin, 20 ng/ml EGF, 100 ng/ml cholera toxin, 50 U/ml penicillin and 50 U/ml streptomycin. In other experiments where the starting ERK needed to be maintained at a low level, the EGF-free recording medium was supplemented with 0.5 mg/ml hydrocortisone, 100 ng/ml cholera toxin, 50 U/ml penicillin and 50 U/ml streptomycin. The cells were incubated in the recording medium for 4 hours before the AC EF simulation.

### EF stimulation

Electric pulses of different waveforms were generated by a programmable function generator (AFG 31000 series, Tektronix). The signal was distributed to a pair of microelectrodes that were 200 µm apart, through a relay matrix controlled by an Arduino board. Isolated amplifiers (Analog Devices, AD215) are installed in the signal path to completely isolate the microelectrodes from all other circuits as shown in Figure 1D. Cyclic Voltammetry (CV) was performed in Fe(CN)_6_^3-^/Fe(CN)_6_^4-^ solution with the microelectrodes, such that the safe range of potential that can be applied was identified where no Faradaic current was detected. The amplitude of all the electric pulses was limited within this safe range to make sure that the signals were only capacitively coupled with no DC components. Cells within 100 µm from the microelectrodes were used to analyze the ERK response by calculating the ERKTR ratio. We define the “threshold” voltage as the minimal amplitude of 50 kHz bipolar square electric pulses that can trigger clear ERK activation, which was the standard waveform of EF stimulation used in previous studies ^43^. Different waveforms of the electric pulses were compared with the same amplitude in each group of experiments. All the following results were reproduced using different microelectrodes with bare Au surface (uncoated) and with HfO_2_ coating of thicknesses ranging from 5 nm to 100 nm. The threshold voltage was about 0.75 to 1V for bare Au surface and ranged from 1.5 V to 8 V for 5 nm to 100 nm HfO_2_ coating, all within the safe potential range verified by cyclic voltammetry scans. The increase of the threshold voltage can be modeled with a simple RC circuit (**Fig. S1**), suggesting that the actual threshold potential drop within the electrolyte was constant and the increase in the total voltage applied is a result of the increased capacitive impedance of the microelectrodes. For clarity and consistency, in the following study, we calibrated the threshold voltage for the actual HfO_2_ thickness on the chip in separate control experiments, use that value for the amplitude of the EF stimulation, and focus on investigating the effect of the waveform and timing characteristics of the applied EF pulses.

### Data processing

During the stimulation and recording experiments, the positions and shapes of the cell nuclei could change with time. To accurately track the cells and identify the fluorescence intensity transfer of the ERKTR-mCherry across nuclei boundaries, we used the Artificial Intelligence (AI) algorithm for segmentation (Segment.ai) and the General Analysis 3 module (GA3) provided by the NIS-Elements (Nikon) for image processing. Specifically, the Segment.ai module was trained with the YFP channel data from EKAR3 to correctly identify all the nuclei boundaries. Then an outer ring-shaped region of interest (ROI) and an inner ring-shaped ROI were generated as the cytosolic ROI and the nuclear ROI for each cell, respectively, by dilation and erosion operations of the boundaries (**Fig. S2**). Within these cytosolic ROIs and nuclear ROIs, the area-averaged intensity of the mCherry channel (from ERKTR-mCherry) was extracted to calculate the ratio of cytosolic to nuclear intensity (ERKTR Ratio Fc/Fn). The data from NIS.ai Module was then exported to Igor Pro (WaveMetrics) for further cell tracking and analysis with a customized script.

## EXPERIMENT RESULTS

### AC EF induces both ERK inhibition and activation with clear waveform dependency

Previous studies on EF modulation of ERK signaling pathway have focused on the activation of ERK, and typically performed in the EGF-free medium where cells are “starved” for 12∼24 hours such that the ERK started at a low baseline. For cells in such an EGF-starved state, after a minimum AC EF stimulation of 1 minute with 50 kHz bipolar square pulses, the ERK level typically started to increase from the baseline in ∼5 minutes and peaked in 10-20 minutes ^43^. Here we performed all experiments in the normal culture medium with the presence of EGF, where the starting ERK level of cells was naturally high, such that we can capture the complete picture of EF-induced ERK dynamics under normal physiological conditions.

For each group of experiments, the minimal amplitude for EF stimulation to trigger ERK activation, i.e., the threshold, was first determined using the 50 kHz bipolar square pulse as the standard waveform (see Methods). At this threshold EF stimulation, a decrease in ERKTR ratio could be first observed, indicating the inhibition of ERK, followed by a pronounced ERK activation peak after 15-20 minutes (Fig.2A). We note that the later ERK activation peak had a consistent time and shape as reported previously ^43^, but the initial rapid ERK inhibition response was unexpected which appeared almost immediately after stimulation (within our time resolution of 30 seconds between images). We then fixed the amplitude, period, and duration of the EF stimulation, and only changed the waveform from square wave to sine, triangular, and sawtooth waves, respectively, to stimulate the same group of cells. Between each trial of stimulation, the cells were given a rest period for 40-60 minutes, to ensure that the signals from one stimulation were not affected by the previous one. At the end of the recording, the square wave was used again to confirm that the response was reproduced from the same waveform and that there was no degradation of cell quality during the whole recording process. Interestingly, responses from the same cells showed clear waveform dependency: the ERK activation peak disappeared when sine (Fig. 2B) and triangular EF (Fig. 2C) were applied but could be recovered with sawtooth EF (Fig. 2D).

**Figure 2.**
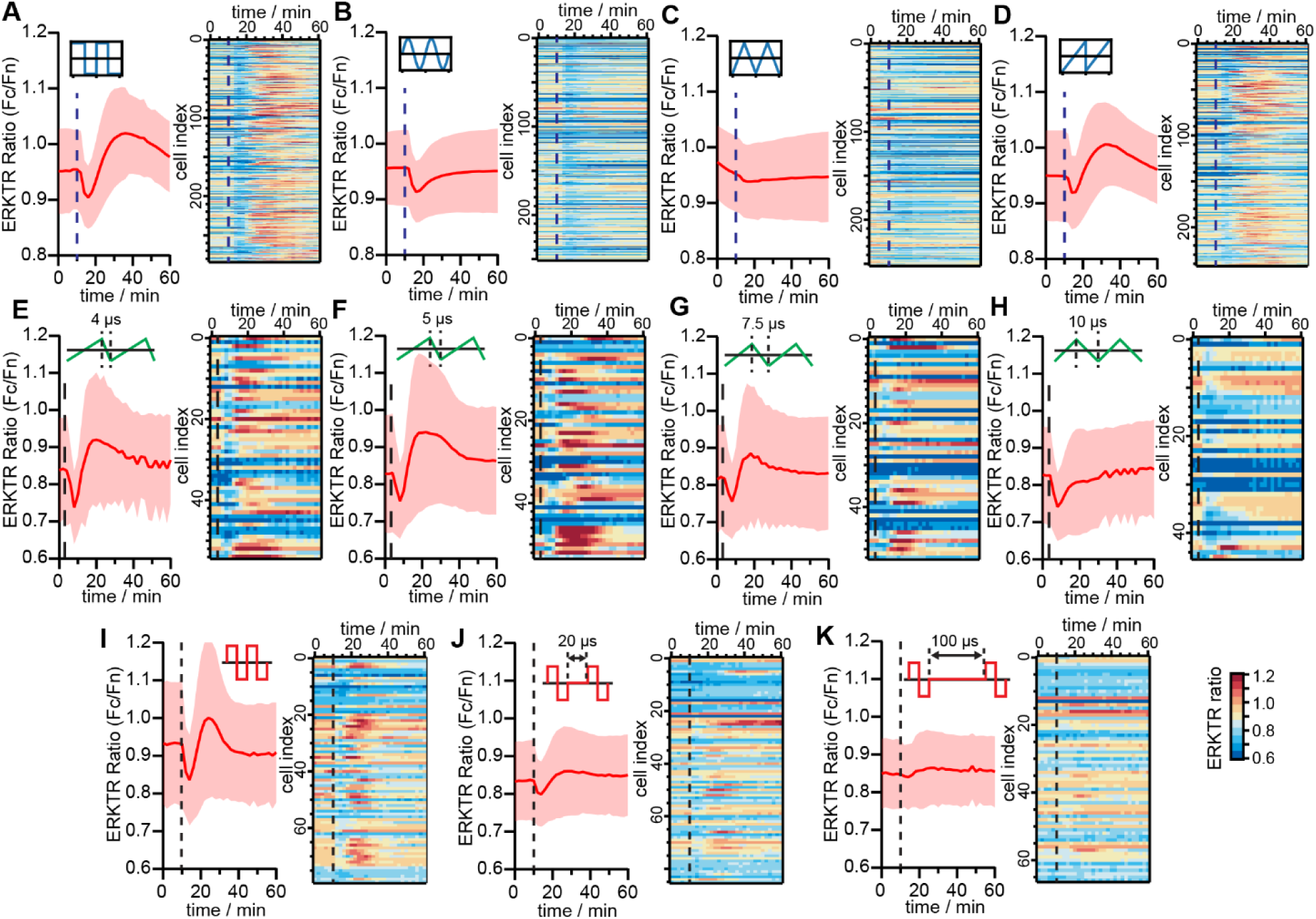
Waveform and timing dependency of EF-induced ERK inhibition and activation. **(A-D)** ERK activities when square, sine, triangle, and sawtooth EF stimulations of the same amplitude and period were applied to the same group of cells. **(E-H)** ERK activities when triangle waveforms of different duty cycles with ramping time from 4 µs to 10 µs applied. **(I-K)** Response of ERK when interval time between individual pulses changed from 0 to 100 µs. The dash lines mark the starting time of the EF stimulations, and the duration of EF stimulations was 3 minutes.

Since all the stimulation parameters except the waveform shape were the same, the data suggested that the ERK activation requires the fast-rising/dropping edges of the signals sent through the microelectrodes, as was only present in the square and the sawtooth. Therefore, we further examined the impact of the ramping speed of the EF on the ERK responses. Namely, we gradually changed the duty cycle of the sawtooth waveform from 99% to 50% to bring the time of the dropping edge from 4 µs (bandwidth limited) to 10 µs for the same amplitude (Fig. 2E-H). The ERK activation peak became weaker and disappeared when the ramping time was longer than 7.5 µs. Notably, the inhibition response was persistent in all cases. In addition, if a rest time of 20 µs to 100 µs was inserted between the 20 µs bipolar square pulses, both activation and inhibition responses significantly decreased (Fig. 2I-K). Overall, we conclude that the inhibition of ERK can be triggered by a broader range of waveforms in comparison to the activation response.

The sensitivity of the ERK signals to the shape of the waveforms and the timing between pulses suggested that the response was not likely attributable to electrochemical processes or chemical species/cues caused by the application of EF in the culture medium, because the range of electrochemical potential sweep at the electrodes was all the same in these tests, and the thin high-k dielectric coating, which further blocked all Faradaic current, did not arrest the response. Last, since these bipolar stimulations did not have a DC component passing through the microelectrodes, there was no net current or ion flow that would impact the cells either. The observed coupling between the AC EF and the ERK signaling should be considered as the electrostatic modulation from the AC components.

### EF-induced ERK inhibition shows a lower threshold and different temporal characteristics than EF-induced activation

The AC EF stimulation by microelectrodes is highly localized. Only cells within 100 µm from the electrodes demonstrate ERK activation following the inhibition period, which is consistent with previous studies ^43^. In contrast, the observed ERK inhibition response to EF stimulation can be observed from cells more than 200 µm away from the electrodes. In addition, at a lower amplitude (∼75% of the threshold required for ERK activation), the EF could be tuned to just trigger the inhibition without activation of ERK for all adjacent cells (Fig. S3). Overall, the inhibition of ERK needed a lower threshold compared to the activation process. Given that the timing and duration of the ERK inhibition are also significantly different from those of the activation response, we further investigated the characteristics of only the isolated inhibition phase without triggering ERK activation by EF.

First, we used “sub-threshold” EF stimulation (at 75% threshold amplitude) to induce only ERK inhibition response from all cells, and gradually changed the duration of stimulation to compare the magnitude of the induced inhibition and their “peak time”, which is the interval between the start of the EF stimulation and the time of the lowest ERKTR ratio. The cells were allowed to rest for 60 minutes after each stimulation before repeating the experiment. The data showed that the ERK inhibition could emerge at a minimum EF stimulation duration of just ∼15 seconds, reaching a consistent peak time of ∼2 minutes as the duration increased. In addition, the inhibition magnitude saturated after ∼ 3 minutes of EF stimulation (Fig. 3B). These characteristics are different from the activation phase, where the minimal stimulation duration required is ∼1 minute, and the peak time is 10-20 minutes ^43^.

**Figure 3.**
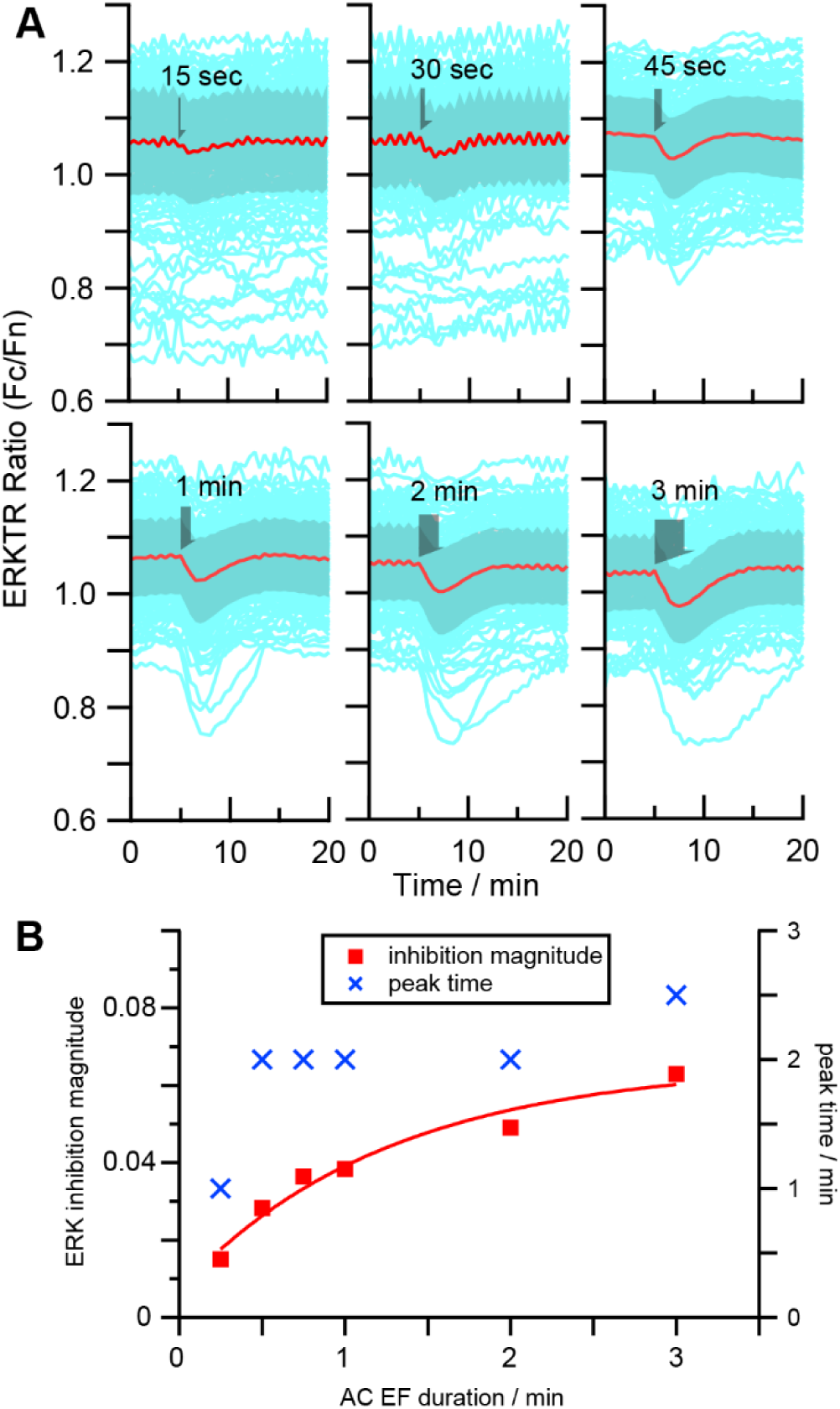
The magnitude and peak time of inhibition depends on the duration of the EF stimulation. **(A)** the ERKTR ratio traces vs time with different durations of EF stimulation from 15 sec to 3 min. The light blue traces are the raw ERKTR ratio from individual cells. The red trace and the shadow denote mean ± standard deviation. **(B)** The ERK inhibition magnitude (red square) and peak time (blue crosses) vs duration of EF stimulation of the same magnitude from the same group of cells, as measured from the average traces (red) of (A).

Second, we scheduled repeated 1-minute-long EF stimulations using bipolar 50 kHz square pulses with interval time gradually decreased from 30 min to 5 min, and the ERK responses from the same group of cells are shown in Fig. 4A-D. The magnitudes of the inhibition extracted from the averaged ERKTR ratio traces are summarized in Fig. 4E. The EF-induced ERK inhibition remained stable for intervals of 30, 20, and 10 minutes, but showed significant decay when stimulations are only 5 minutes apart, giving a refractory time for EF-induced inhibition between 5-10 minutes. As a comparison, EF-induced activation is known to have a much longer refractory time >20 minutes ^44^.

**Figure 4.**
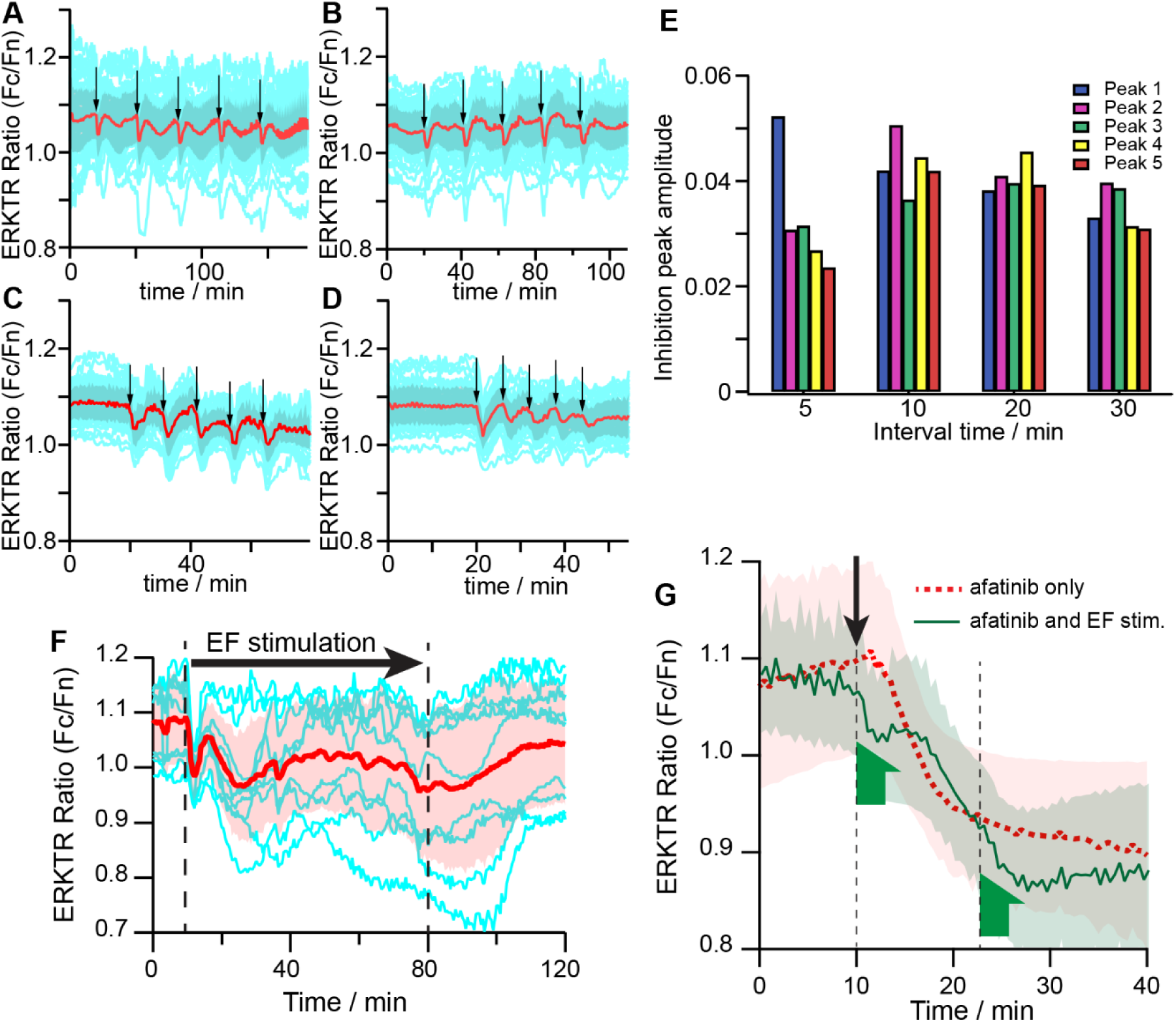
Time characteristics of EF-induced inhibition and comparison with EGFR inhibitor. (A-D) Periodic modulation of ERK inhibition induced by short EF stimuli. Representative time traces of ERKTR ratio of individual cells when multiple cycles of 1 min EF stimuli are applied to the same group of cells with different time intervals, thicker lines are population average, shadow region shows 25^th^ and 75^th^ percentile values. Interval time: (A) 30 min, (B) 20 min, (C) 10 min, (D) 5 min. (E) Comparison of ERK inhibition peak amplitude of different interval times in (A-D). (F) Fluctuation of ERK with inhibitions when long sub-threshold 50kHz square wave EF stimulation was applied. Dash lines mark the beginning (at 11 min) and the end (at 80 min) of the EF stimulation. (G) Red trace: ERKTR ratio showing ERK inhibition induced when 1 µM EGFR inhibitor afatinib was applied (marked by the black arrow). Green trace: ERKTR ratio showing inhibitive ERK response as two separate 3 min long EF stimulations (marked by the green half arrows) were applied after the application of afatinib (marked by the black arrow).

Third, we performed a sustained sub-threshold EF stimulation for an extended hour. After the rapid inhibitive ERK peak in the first 10 minutes, a slow fluctuation of ERK level that was overall below the baseline could be observed (Fig. 4F). Comparing the two inhibitive responses, the earlier rapid peaks were well aligned and consistent between cells as have been shown previously, while the later slow fluctuations were not as well synchronized and demonstrated larger variations in terms of magnitude and period from cell to cell. In addition, after the EF stimulation was stopped, the ERK gradually returned to the normal level in about 30 minutes, which showed that the lower ERK level was not due to photobleaching or cell degradation. The initial fast and later slow fluctuations of inhibitive ERK responses during the EF stimulation suggested that there may be several signaling feedback processes of different time scales that are involved in generating this rich dynamic behavior.

The energy from the AC EF stimulation <10 MHz is mostly coupled with constituents in the cell membrane region ^42, 49^. A previous study has shown that EF-induced phosphorylation of EGFR can lead to the activation of ERK ^43^. Are these inhibitive effects also tied to EF-induced changes in EGFR? To answer this question, we compared the ERK inhibition induced by EF and chemical methods. Specifically, we first added 1 µM of afatinib to the culture medium to inhibit the phosphorylation of EGFR and recorded the time course of the ERK decrease as the control experiment (Fig. 4G, red trace). Then, in a separate run, a 3-minute of sub-threshold EF stimulation was applied right after adding the same amount of afatinib in the medium. The ERK level rapidly decreased to a deeper inhibition state during this period, but when the EF stimulation was stopped, it recovered briefly and returned to follow the same trend of slower decrease as in the control experiment (Fig. 4G, green trace). At around 22 minutes when the ERK reached the same level as in the control group, another 3-minute long EF stimulation was applied, and again a deeper inhibition state showed up, which recovered afterward to the same plateau after EF stimulation stopped. The much more rapid time characteristics of EF-induced inhibition and its relative independence from the chemical EGFR inhibitor suggested that the EGFR may not be the only component impacted by EF at the cell membrane.

### EF stimulation may perturb the activity of Ras at the cell membrane

The lipid-anchored Ras-GTP is activated by the Grb2/Sos after EGFR phosphorylation. It serves as the amplifier to regulate downstream ERK signals (Fig. 1A). If the activity of the Ras-GTP complex were also influenced by the EF stimulation, it would likely cause a more rapid impact in the downstream signaling. Here we evaluate this possibility by quantifying the EF-induced changes in the Ras-GTP at the membrane. Specifically, we mixed an equal amount of the MCF10A-EKAR3-ERKTR cells with the MCF10A-FRB-RBD cells and co-cultured them on the same chip. The MCF10A-EKAR3-ERKTR cells give the ERK response in the RFP channel as in previous experiments, and the MCF10A-FRB-RBD cells have been transfected with Lyn11-FRB-CFP, which is used to mark the position of the lipid membrane, and RBD-GFP, which can track the activated Ras-GTP (Fig. 5A).

**Figure 5.**
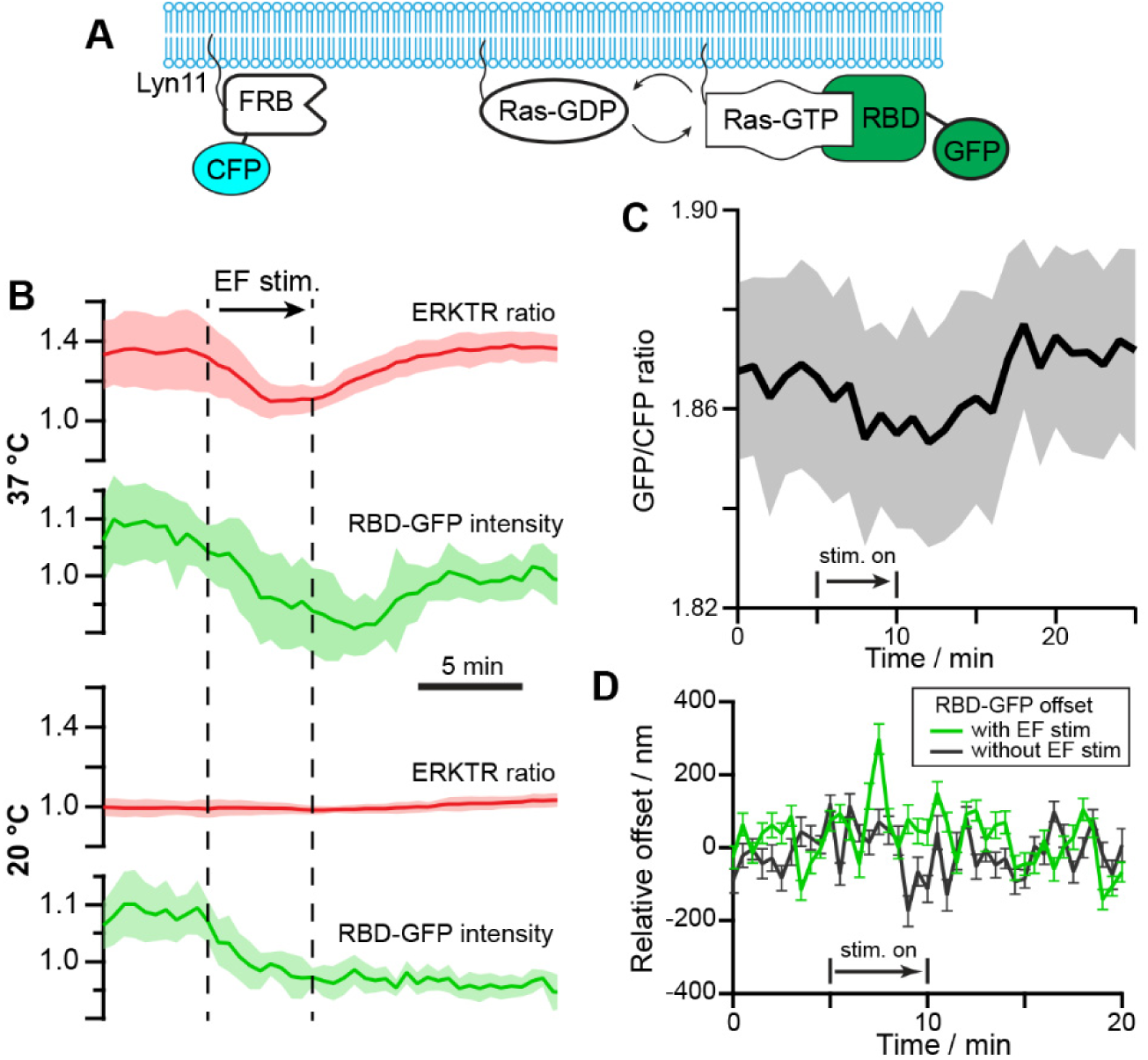
Ras-GTP changes during EF stimulation. (A) Schematics of the Lyn11-FRB-CFP and RBD-GFP reporter expressed in the MCF10A cell line. (B) Comparison of ERK and Ras response at different temperatures with EF stimulation. Top group: at 37 °C the ERK showed clear inhibition followed by recovery to activation upon EF stimulation, and the RBD-GFP intensity showed a correlated decrease followed by recovery within the same period. Bottom group: at 20 °C, the ERK level of the same cells remained *at a low level* during EF stimulation, and RBD-GRP intensity only showed an immediate decrease during EF stimulation without recovery. The thick lines give the average results from cells (n=6) with standard deviation shown as the shadowed area. (C) The ratio of fluorescence intensity from RBD-GFP and Lyn11-FRB-CFP showed a clear decrease and recovery during and after EF stimulation. The thick line gives the average results from different cells (n=6) with standard deviation shown as the shadowed area. (D) The offset of the peak positions of RBD-GFP and Lyn11-FRB-CFP with and without EF stimulation. The peak position was calculated as a Gaussian fit in a line profile crossing the cell membrane in the fluorescence images. The chromatic offset between GFP and CFP has been subtracted from the data.

With a spinning disk module (BD CARVII) we captured the confocal images at the basal membrane of the MCF10A-FRB-RBD cells to obtain the average intensity of RBD-GFP. The changes in RBD-GFP during EF stimulation was then compared to the ERKTR-ratio from the RFP channel signals of the co-cultured MCF10A-EKAR3-ERKTR cells in the same images (Fig. 5B). At normal cell culture temperature (37 °C), a rapid decrease of the RBD-GFP intensity was recorded right after the EF stimulation started, which synchronized with the initial inhibition phase of ERK. After about 10 minutes, the RBD-GFP intensity returned to the original level, and the ERKTR-ratio turned back high during the same period. As a control, the temperature of the environmental chamber was cooled to room temperature at 20 °C, at which cell activities slowed down and the ERK level of the cells all switched to low gradually. Upon EF stimulation, the same decrease of RBD-GFP can be seen, however, with no recovery afterward, and the ERK level stayed low with no response to the EF at all.

To minimize the impact of photobleaching in the analysis, we further used the Lyn11-FRB-CFP as the internal calibration to quantify the changes in RBD-GFP. Since no Rapamycin was introduced to our system to activate FRB, the intensity of the CFP signals could serve just as the baseline to compare with other membrane-anchored molecules such as activated Ras. We calculated the intensity ratio of the GFP and CFP signals at the cell membrane from different cells, and showed that the GFP/CFP ratio gave the same rapid decrease after EF stimulation, followed by the recovery to a higher level in ∼10 minutes (Fig. 5C). The consistent results in Fig. 5B and 5C suggested that there was an EF-induced change of Ras-GTP density at the cell membrane, and its temporal characteristics were aligned with the observed ERK inhibition and activation phases. The initial disruption of activated Ras seemed to be a direct effect of the EF stimulation, and its later recovery to a high state was correlated to the signaling activities associated with the activation of ERK.

An interesting question is: could the EF impact on the Ras activities be associated with a net force exerted on the membrane protein? If so, this could show up in the relative position change of the RBD-GFP molecules from the membrane. We set the focal plane at 4 µm above the coverslip to get the cross-section confocal images of the cells. The intensity line profiles crossing the cell membrane at different positions were extracted from the GFP and CFP channels and fitted with Gauss functions (Fig. S4). The offset between the peak positions of the GFP and the CFP signals was used to evaluate how the GFP-labelled Ras shifted relative to the cell membrane. We note that the applied EF pulses were bipolar, and has a much shorter period (20 µs square wave) than the exposure time (500 ms). As a result, the force felt by the molecules could be symmetric about the membrane and we would not see any net position change. Or these images did not have enough time resolution to capture the real-time response of the GFP-labelled molecules to the fast-alternating EF perturbations, but could only produce the average offset as a “residue effect” between each exposure. We saw no significant offset changes between the GFP and CFP signals in most of the line profiles before/during/after the EF stimulation, which means that on average the RBD-GFP appeared to follow the cell membrane consistently. Only in a small portion of the line profiles data (∼10%), a large discrepancy between the GFP and CFP peak positions could be seen just during the EF stimulation (Fig. 5D), capturing possibly Ras detaching events due to EF perturbation. We note that these events were only captured in a small subset of data, and more direct evidence and quantification with higher time resolution would be needed to better track the real-time response of membrane molecules to EF. For example, nanometer-level membrane deformations can be detected using differential detection method ^50^, which could track the fluctuation of the surface tension/energy of the cell membrane associated with the ligand-binding events and membrane protein concentration changes. In addition, the electric potential fluctuations propagating along the membrane of neuron cells could also be detected by such mechanically amplified deformation with high temporal resolution ^51^. Therefore, further investigations with faster imaging techniques and high-resolution analysis methods could shed more light on the direct biophysical impact of EF stimulation on membrane proteins.

## DISCUSSION

### Energy from AC EF can couple with the cell membrane constituents

Overall, two main observations suggest that the coupling of the EF energy to the membrane region is dissipative through electrostatic interactions: (1) The ERK response was sensitive to the waveform and timing of EF pulses when all other parameters were the same and Faradaic process was absent; (2) A minimal amplitude and duration of stimulation was required to produce an appreciable effect.

Electric fields of different frequencies influence the cell in distinct ways. A DC EF can introduce electrolytic effects or ion-concentration polarization. But there is overall little biophysical effect at the subcellular level to DC EFs other than the transient discharge current across the membrane ^49^. On the other hand, AC EF components exhibit two effects: conductive power dissipation, caused by the translational friction of current carriers (ions), and dielectric relaxation, associated with rotation or flexion of molecular dipoles. It has been shown that the power dissipation in the cell membrane region starts to increase when the frequency of the AC EF is higher than ∼100 kHz, and significantly exceeds the external medium in the MHz and lower GHz range’ due to the increasingly dominant contribution from the dielectric relaxation component. The increase of the energy coupling within this frequency range occurs mainly because the head groups of the membrane lipids have limited mobility within the ordered membrane structure. Thus, even in EF exposures that do not cause a significant energy increase at the macroscopic whole-system level in the form of temperature rise, the locally increased power dissipation in cell membranes can lead to significant consequences at the microscopic single-cell level ^49^. We may borrow a few ideas from the recent studies of the nonlinear dielectric spectroscopy ^52^, and consider the cell membrane as a heterogeneous nanothermodynamics system of an ensemble of independently relaxing “regions” ^53, 54^. Different degrees of freedom are weakly coupled to each other while all connected to a common thermal reservoir. Such a heterogeneous system can demonstrate energy absorption through dielectric relaxation with characteristic frequency^52^ that changes “fictive” temperature of local degrees of freedom and drive the activities of membrane constituents.

It should be noted that the models of calculation have been based on a simple lipid bilayer structure with no other membrane components ^49^. Therefore, for real cells the effective frequency where EF energy couples with different membrane constituents such as large membrane proteins/complexes can extend to a much broader range. Although the time scale associated with the motions of molecular changes is not known, NMR ^55, 56^, fast laser spectroscopy ^57^, and MD simulations in other protein systems ^58^, broadly suggest time scales on the order of 1-100 µs for typical macromolecular transitions, corresponding to 10KHz ∼ MHz range, which coincides with our current experiments.

### Membrane proteins as sensors of EF perturbation

Apart from dielectric relaxation, EF-induced transmembrane potential fluctuations could also affect the conformational changes of membrane proteins, although the AC EF applied in our studies is rather weak (between 10∼100 V/cm) as compared to most models or what is used in electroporation experiments. Recent studies have shown that the farther the membrane protein extends away from the cell membrane, and the less rigid it is, the more strongly the protein could respond to perturbations in the transmembrane potential ^59^. Many key membrane proteins and protein complexes in the ERK signaling pathways, including EGFR and Ras, extend several Debye lengths away from the membrane and contain flexible, charged/polarized subdomains. Moreover, it has been shown that the electrostatic interaction between the EGFR and membrane lipids microdomain/rafts may also play an important role in the autoinhibition and activation process ^60^. Therefore, membrane protein-initiated activities could in principle be sensitive to electric perturbations.

Moreover, we have seen that EF perturbation can cause both inhibition and activation of the ERK signaling pathway, with very different time characteristics. While the onset time of the EF-induced ERK activation was consistent with those observed under chemical stimulation ^46^, the EF-induced ERK inhibition effect was faster than that from the chemical EGFR inhibitor. Our data strongly suggest that EGFR is probably not the only membrane component responsible for the bipolar ERK responses to EF stimulations.

### Possible mechanism of the inhibitive ERK response

The ERK signaling pathway is regulated by many negative feedback loops, including direct phosphorylation by ERK and RSK2, and the transcriptionally induced feedback regulators ^61^. However, it doesn’t seem likely that the inhibitive ERK response to EF can be directly attributed to these feedback loops because it takes typically ∼10 minutes for an appreciable increase in ERK to occur first. Instead, the rapid inhibition and an overall bipolar ERK response to EF suggested that an additional cell membrane component downstream to EGFR was involved, and its effect on ERK is more immediate. A possible candidate is the Ras protein and its active membrane-anchored complex formed during the signaling process, whose activity depends on the interaction with the cell membrane and clusterization.^62, 63^ We have observed the synchronized reduction in the active Ras during the inhibitive phase of the ERK change, followed by the correlated recovery in both signals. Although correlation doesn’t imply causality, it is reasonable to speculate that the change in Ras could contribute to the initial rapid ERK inhibition, because its function as the amplifier in the signaling pathway requires forming a membrane-anchored complex, which could be susceptible to the EF perturbation.

### Limitations and prospects

This study focuses mainly on the phenomenological investigation of how ERK signaling responded to different EF parameters. The results showed that the perturbation is electrostatic and mostly consistent with models of energy coupled through frequency-dependent dielectric relaxation. The data also suggested that multiple cell membrane components contributed to the EF-induced bipolar ERK response, including possibly Ras and EGFR, responsible respectively for a rapid inhibition phase and a later slower activation phase with different thresholds and sensitivity to EF waveforms. However, it remains an open question how the EF perturbation specifically changes the states and activities of EGFR and Ras at the molecular level. More direct evidence and modeling are still needed to understand the details. Currently, the fluorescence imaging in our experiments is too slow to capture the real-time perturbation of membrane components, and tracking the conformational states during stimulation is also challenging. We could only observe what is likely the residue effect in terms of concentration and position at a much longer time scale than the EF pulses. A promising development of the differential detection method using fast brightfield phase-contrast imaging, which has demonstrated sub-nm accuracy and sub-ms time resolution ^51^, could be a promising way to capture the real-time membrane response to the fast EF perturbation, although tracking specific species on the membrane would still be a challenge. Overall, our work is a first step in exploring the interesting effect of exogenous AC EF on intracellular signaling pathways. If the molecular mechanism can be further understood and developed as a general method, it could provide a new dimension of parameter space and controllability to precisely modulate specific cell signaling pathways with carefully engineered EF patterns that can achieve high temporal and spatial resolutions, which can be an advantage compared to chemical modulations, such that the behavior and/or fate of targeted cell population can be guided and programmed selectively.

## Conclusion

In summary, we employed dielectric-coated microelectrodes and various reporters to study the ERK signaling pathway in MFC10A cells responsed to different EF stimulations with precise spatial and temporal resolution in a Faradiac process-free environment. Under normal culture medium where EGF was present, we observed an interesting rapid inhibition phase in the ERK signal shortly after EF stimulation with a 50kHz bipolar square wave, followed by an activation phase that was much slower and broader. The inhibition required a lower EF amplitude threshold, shorter stimulation time, and faster refractory time in comparison to the activation response. In addition, both inhibition and activation of ERK demonstrated clear sensitivity to the waveform and timing of the electric pulses used in the EF stimulation. There was a correlated change in the active Ras-GTP to the ERK signal changes during the EF stimulation. We propose that the EF stimulation could couple with the cell membrane through the frequency-sensitive dielectric relaxation process, and perturbed the activities of both Ras and EGFR, which resulted in the bipolar ERK response with a rich time dynamics. If this model can be verified and understood with more direct and quantitative evidence, it can serve as a new way to precisely modulate signaling pathways with exogenous EF.

## ACKNOWLEDGEMENTS

Q.Q. acknowledge the support by the Air Force Office of Scientific Research under award number FA9550-16-1-0052, and DURIP by the Air Force Office of Scientific Research under award number FA9550-21-1-0228. We acknowledge Dr. John Albeck at University of California, Davis, for providing the MCF10A-EKAR3-ERKTR cell lines.

## AUTHOR CONTRIBUTIONS

Q.Q. designed research; M.H. and H.L. performed the experiments; M.H., H.L. and Q.Q. analyzed the data; K.Z., L.G., M.Z., H.Z., P.N.D. discussed the analysis and interpretation; All authors contributed to the preparation of figures and the manuscript.

