## Supplementary Information for "Electrostatic modulation of signaling at cell membrane: Waveform- and time-dependent electric control of ERK dynamics"

**This file includes:**

Supplementary Figures S1 to S4

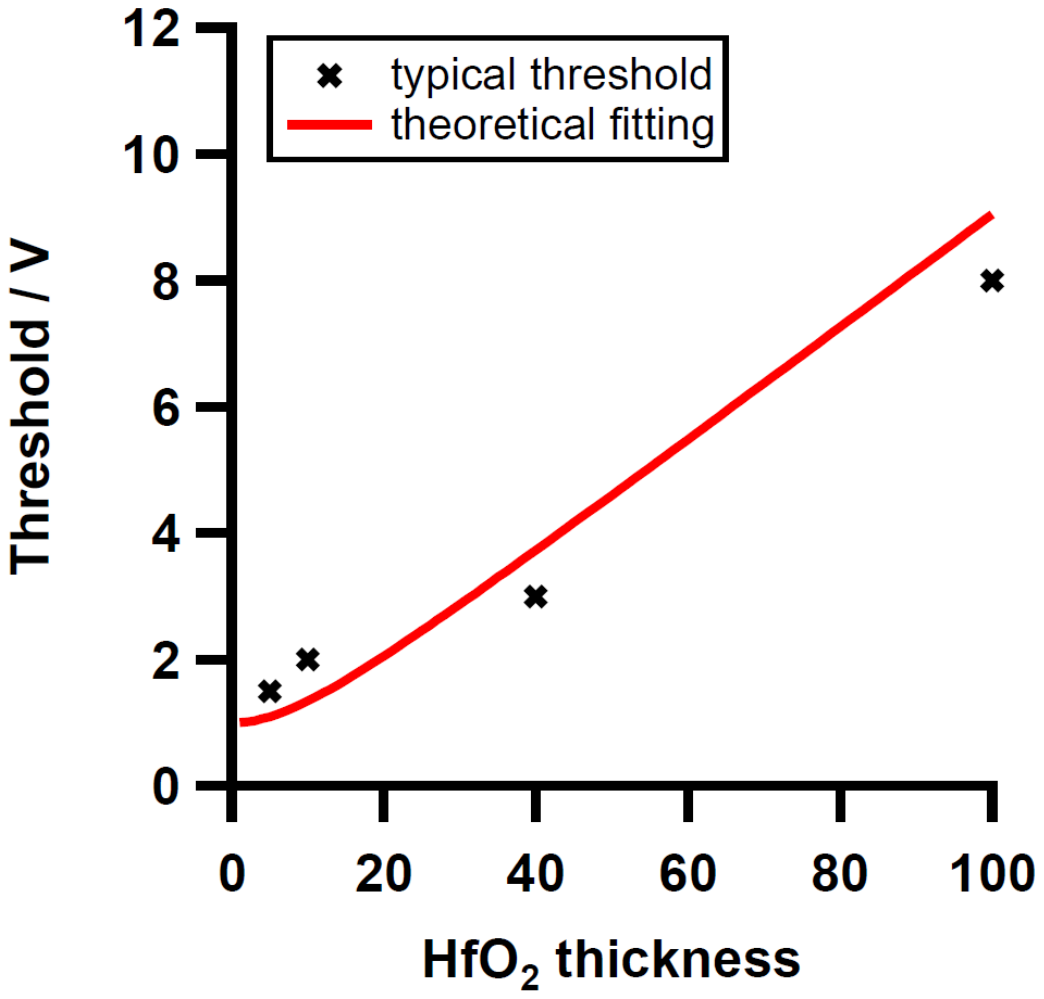

**Fig. S1**. Evaluation of electrode stability. (a) Cyclic voltammetry scan of gold microelectrode in culture media DMEM/F-12(Cat#21041025, Life Technologies) with two electrodes configuration, scan rate 25 mV/s. (b) Optical microscope image of gold electrode before and after bipolar 10µs wide ±1.5V EF pulses were continuously applied for >1 hour.

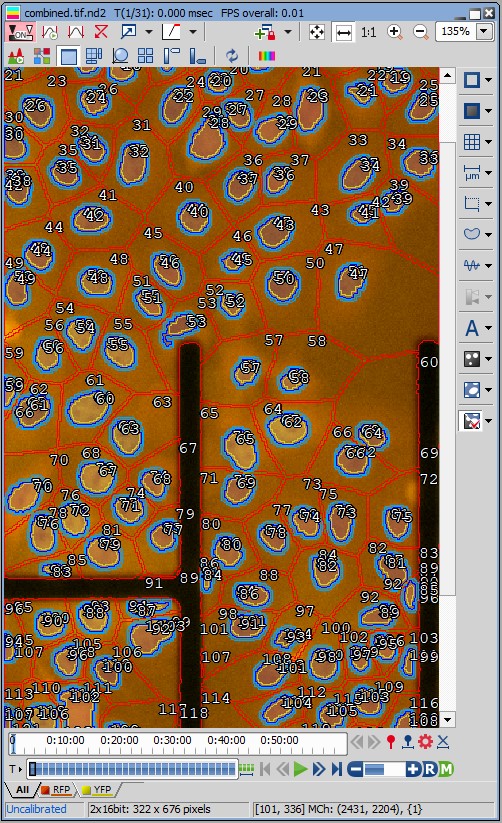

**Fig. S2.** Cell segmentation and automated calculation of ERKTR ratio using NIS-Elements with Segment.ai package. The YFP channel is used for training and identification of cell nucleus boundary (dark blue lines). The average fluorescence intensity within the nucleus region (light yellow) and the average fluorescence intensity of the surrounding region in the cytoplasm (light blue belt) are calculated from the RFP channel, and used for the calculation of ERKTR ratio calculation.

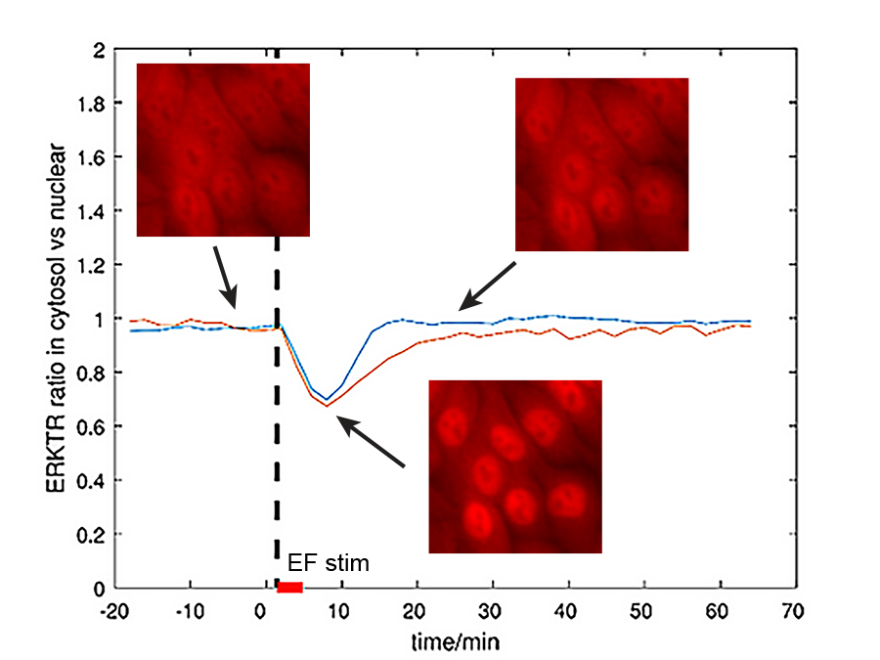

**Fig. S3.** EF can induce rapid inhibition only without causing a later activation at a lower amplitude.

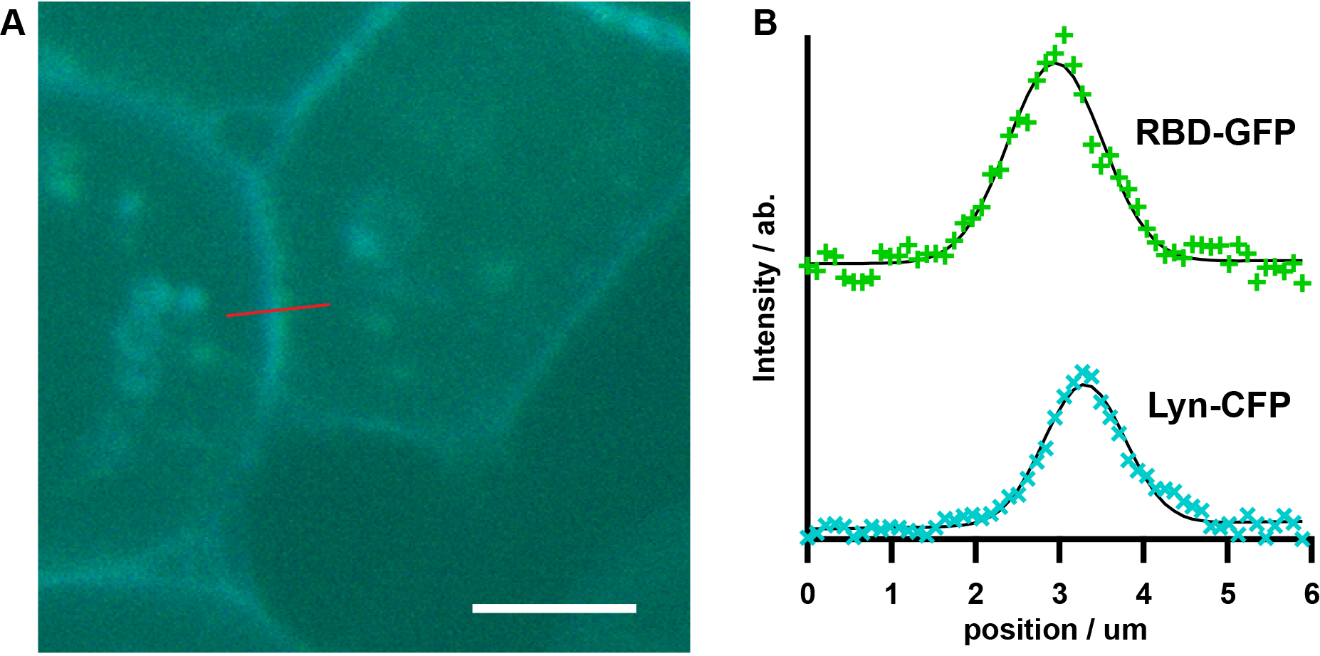

**Fig. S4.** The line profile analysis of the Raf-RBD-GFP and Lyn-FRB-CFP channels. The intensity profiles are extracted from the image stack and fitted by Gaussian functions. The peak positions of the two channels are used to calculate the relative offset of the Ras-RBD-GFP and Lyn-FRB-CFP molecules. The Lyn-FRB-CFP is used as the cell membrane position reference to track the relative motion of the Ras-RBD-GFP molecules under EF modulation. The chromatic offset error between the two channels is not corrected but since we only use the relative changes, the absolution offset does not matter.
